# Glucan unmasking identifies regulators of temperature-induced translatome reprogramming in *C. neoformans*

**DOI:** 10.1101/2020.12.15.422993

**Authors:** Amanda L.M. Bloom, David Goich, Corey M. Knowles, John C. Panepinto

**Affiliations:** Department of Microbiology and Immunology, Witebsky Center for Microbial Pathogenesis and Immunology, Jacobs School of Medicine and Biomedical Sciences, University at Buffalo, SUNY, Buffalo, NY 14203, USA

## Abstract

The cell walls of fungi are critical for cellular structure and rigidity, but also serve as a major communicator to alert the cell of the changing environment. In response to stresses encountered in human hosts pathogenic fungi remodel their cell walls. Masking the b-1,3-glucan component of the cell wall is critical to escape detection by innate immune cells. We previously demonstrated that b-1,3-glucan is unmasked in response to host temperature stress when translatome reprogramming is defective in *C. neoformans*. Here, we used b-1,3-glucan unmasking as an output to identify signaling modules involved both in masking and translatome reprogramming in response to host temperature stress. We reveal that the High Osmolarity Glycerol (HOG) MAPK pathway is involved in translatome reprogramming and that mutants in this pathway display moderate unmasking when grown at 37°C. Additionally, we show that mutants of the Cell Wall Integrity/Mpk1 MAPK pathway extensively unmask b-1,3-glucan. While the CWI pathway does not impact translatome reprogramming, our data suggest it may play a role in the post-translational regulation of transcription factors that govern masking.

**Importance:** Cryptococcus neoformans is a fungal pathogen that causes devastating morbidity and mortality in immunocompromised individuals. It possesses several virulence factors that aid in its evasion from the host immune system including a large polysaccharide capsule that cloaks the antigenic cell wall. Studies investigating how the cell wall is remodeled to keep this pathogen disguised in response to stress have been limited. We previously found that host temperature stress results in translatome reprogramming that is necessary for keeping the highly antigenic β-(1,3)-glucan component masked. Our data reveals signaling modules that trigger these responses and suggest the points of regulations at which these pathways act in achieving masking. Understanding these mechanisms may allow for therapeutic manipulation that could promote immune recognition and clearance of this fungal pathogen.

## Introduction

*Cryptococcus neoformans* is an environmental basidiomycete fungus that is able to enter the mammalian lung and persist. In cases of defective adaptive immunity, it can cause pulmonary infection, or disseminate to the central nervous system where is causes a deadly meningitis. The primary comorbidity associated with cryptococcosis is HIV co-infection, however recent use of anti-TNF-a monoclonal antibodies in the context of rheumatoid arthritis is associated with increased risk of cryptococcosis (Liao et al., 2016).

*C. neoformans* possesses multiple complex traits that promote its persistence in the lung and its virulence. One of these traits, thermotolerance, is required for mammalian infection. Previous work in our laboratory has implicated deadenylation-dependent mRNA decay mediated by Ccr4 in promoting adaptation to host temperature (Havel et al., 2011; Bloom et al., 2013; Bloom et al., 2019). This work suggests that reprogramming of the mRNAs associated with translating ribosomes, the translatome, is a prerequisite for temperature adaptation (Bloom et al., 2019). Coupled to temperature adaptation, is the maintenance of cell wall integrity, and the masking of the ubiquitous fungal pathogen-associated molecular pattern (PAMP), b-1,3-glucan.

Another environmental *Cryptococcus* species, *C. amylolentus*, is deficient in thermotolerance, and like our *ccr4*D mutant, also unmasks b-1,3-glucan at 37°C (Bloom et al., 2019). In addition, *C. amylolentus* fails to repress abundant ribosomal protein (RP) transcripts, which is necessary for reprogramming the translatome in response to temperature stress. This led us to a model in which thermotolerance and b-1,3-glucan masking are linked, likely via changes that ensue from translatome reprogramming.

We’ve previously coined the term “adaptive agility”, defined as the extent and speed at which an organism can reshape its proteome to one that is suited for its new environment, and speculated that it is essential for and determines an organism’s pathogenic potential (Leipheimer et al., 2019). Host-temperature induced mRNA decay and translatome reprogramming occur immediately in response to stress (Bloom et al., 2019). Post-translational modifications of proteins allow for rapid, efficient, and dynamic responses by altering the function of proteins that are already present (Zhang et al., 2015; Spoel, 2018). There have been several examples of how the translational landscape of fungi is regulated by kinases in response to different stressors (Leipheimer et al., 2019).

The b-1,3-glucan component of the fungal cell wall is highly antigenic. The pattern recognition receptor Dectin-1 specifically recognizes b-1,3-glucan and promotes phagocytosis and inflammatory responses (Brown et al., 2003). In response to several stressors encountered in the human host, *Candida albicans* engages specific pathways that reduce the exposure of this PAMP. The CEK1-mediated MAPK pathway seems to generally regulate b-1,3-glucan masking (Galan-Diez et al., 2010), signaling through the protein kinase A pathway is required for b-1,3-glucan masking in hypoxic and iron limiting conditions (Pradhan et al., 2018; Pradhan et al., 2019). Interestingly, the Gpr1 receptor, but not the canonical downstream cAMP pathway, regulates lactate-induced masking in *C. albicans* (Ballou et al., 2016). While the cell wall integrity MAPK pathway has been identified in *C. neoformans* (Gerik et al., 2005), pathways that regulate glucan masking specifically in response to stressors have not been investigated.

In this report, we investigate the linkage between glucan masking, translatome reprogramming and thermotolerance by using b-1,3-glucan unmasking as the phenotypic output of a screen of kinase deletion mutants. We identify two core MAP Kinase modules that mediate glucan masking. Mechanistic investigation of point of action suggests that the Hog1 pathway regulates unmasking via translatome reprogramming, whereas the cell integrity pathway likely regulates unmasking post-translationally, perhaps through the activation of transcription factors.

## Materials and Methods

### Strains used in this study

For all experiments the *C. neoformans* serotype A *var. grubii* strain H99 was used as WT. Kinase mutants were created by the Bahn laboratory (Lee et al., 2016) and obtained from the fungal genetics stock center. The *ccr4*Δ mutant was created by biolistic transformation of the H99 strain using a PCR amplified DNA construct in which the nourseothricin resistance cassette was flanked by sequences upstream and downstream of the *CCR4* gene. The URA resistance cassette was dropped from our previously constructed plasmid (Panepinto et al., 2007) and replaced with NAT. Primers for cloning and amplification are listed in Table 1. The knock out clone was verified by PCR, Southern and northern blot.

**Table 1.**
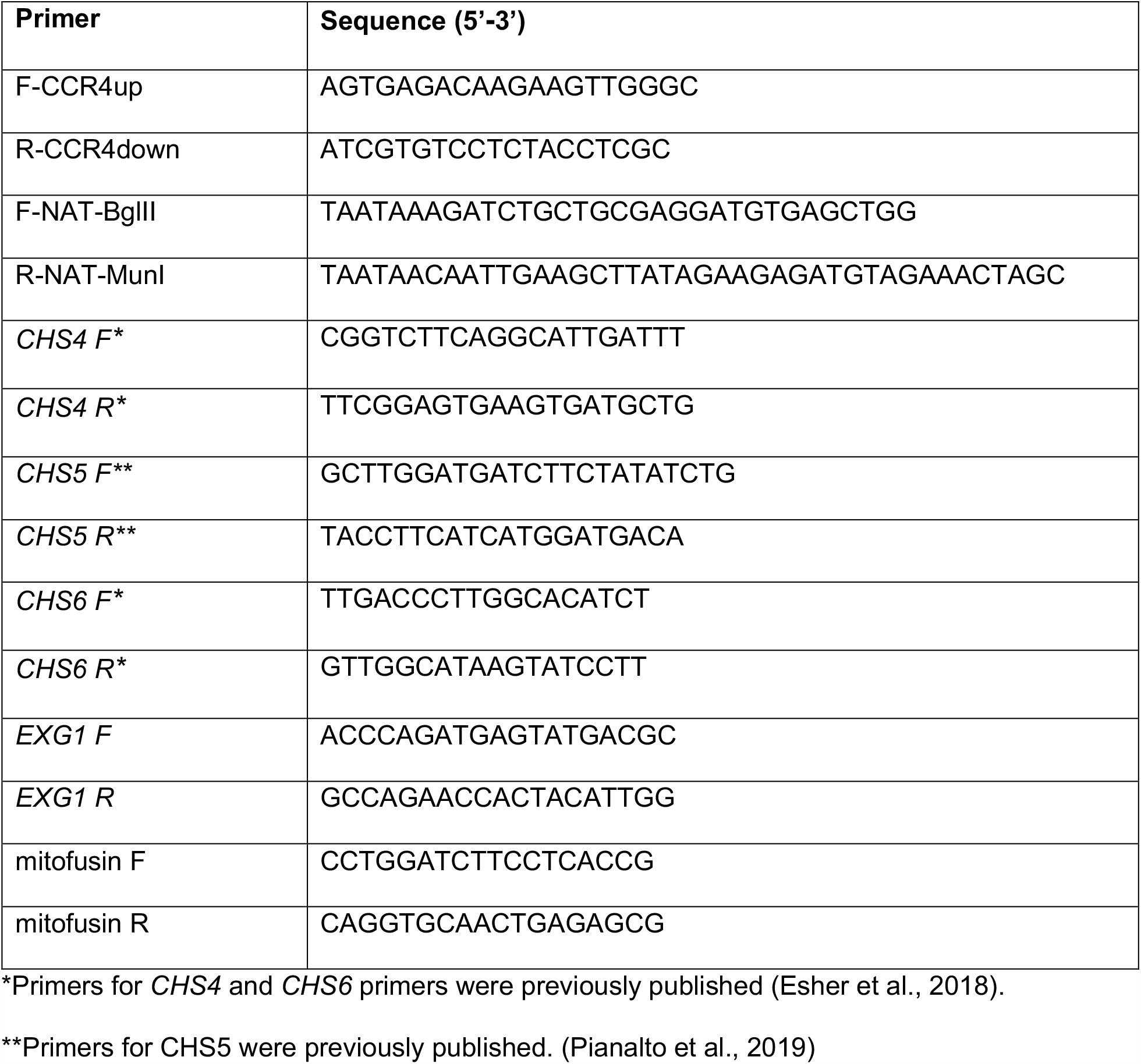
Oligonucleotide sequences used in this study.

### Beta-1,3-glucan staining and detection

For screening of the kinase mutant library, cells were grown overnight in 100 μL of YPD in a 96-well plate at 37°C. Cells were pelleted, washed twice in PBS, and then stained in the 96-well plate following the staining protocol below. For verification of identified mutants from the screen, cells were grown in 3 mL of YPD overnight at 37°C, shaking at 250 rpm. Cells were washed once in PBS and resuspended in PBS to an OD_600_ = 1.0. For each strain, 100 μL of cells was aliquoted to 2 microfuge tubes and pelleted. For staining, cells were resuspended in 100 μL of PBS containing 15 μg/mL anti-β-(1,3)-glucan antibody (Biosupplies Australia). For no primary controls, cells were resuspended in 100 μL of PBS. All tubes were incubated at 37°C with gentle rotation for one hour. Cells were pelleted and washed 3 times with PBS and resuspended in PBS containing 10 μg/mL Alexafluor 488-goat anti-mouse secondary antibody. Cells were incubated at 37°C with gentle rotation for 40 minutes protected from light. Cells were pelleted and washed three times in PBS, then fixed in 3.7% formaldehyde at room temperature for 20 minutes protected from light. Finally, cells were washed and resuspended in PBS. For microscopy cells were visualized and images captured using a Leica SP8 TCS Confocal microscope. For flow cytometry, fluorescence was measured using a BD LSRFortessa flow cytometer. Statistical anlysis for flow cytometry was done using Graphpad Prism (Version 8.4.3).

### Growth curves

All strains were grown overnight in 3 mL of YPD at 30°C, shaking at 250 rpm. Overnight cultures were used to seed fresh YPD cultures to an OD_600_ = 0.15 and were grown at 30°C for 5 hours. These mid-logarithmic cultures were then diluted to OD_600_= 0.1 in 100 μL final volume in a 96 well plate. Growth was assessed by measuring OD_600_ every 10 minutes for 20 hours using a Biotek Synergy Hybrid plate reader.

### Spot plate analyses

All strains were grown in 3 mL of YPD overnight at 30°C. Cells were washed twice with sterile deionized water (SDW), and resuspended in SDW at OD_600_ = 1.0. Five 10-fold serial dilutions were prepared in SDW and 5 μL of each dilution were spotted onto YPD agar plates supplemented with cell wall inhibitors (0.5 mg/mL caffeine, 1 mg/mL calcofluor white, 5 mg/mL congo red, 0.03% SDS). Plates were incubated 30°C or 37°C for 3 days and photographed.

### Polysome profiling

Polysome profiling was done as previously published (Bloom et al., 2019) with minimal differences. Briefly, cells were grown to mid-log phase in YPD and half of the culture was pelleted, resuspended in pre-warmed 37 °C YPD, and incubated at 37 °C for 30 minutes. Cycloheximide (0.1 mg/mL) was added immediately and cells were pelleted by centrifugation at 4000 rpm, 4 °C, for 3 minutes. Pellets were flash frozen in liquid nitrogen and stored at -80°C until lysis. Pellets were washed in polysome lysis buffer (20 mM Tris HCl, pH 8.0, 140 mM KCl, 5 mM MgCl_2_, 1% Triton X-100, 25 mg/mL heparin sodium sulfate, 0.1 mg/mL cycloheximide) and pelleted. Cells were resuspended in residual buffer, transferred to a microfuge tube, and pelleted at 13,000 rpm for 30 seconds. Supernatant was removed, and pellets were resuspended in 100 µl of cold lysis buffer, and transferred to an Eppendorf tube containing glass beads. Cells were lysed mechanically in a Bullet Blender for 5 min followed by addition of 300 µl cold lysis buffer. Supernatant was transferred to a cold microfuge tube and centrifuged for 5 min, at 14,000 rpm, 4 °C. Cleared lysates were quantitated for RNA, and 250 µg was loaded on top of 10–50% sucrose gradients. Gradients were subjected to ultracentrifugation for 2 h, 39,000 rpm, 4 °C. Sucrose gradients were then pushed through a flow cell, and RNA was detected by UV–vis A254 absorbance.

### Northern blot assessment of *RPL2* mRNA repression

For all strains, cells were grown in 3 mL of YPD at 30°C, 250 rpm overnight. Overnight cultures were used to seed 40 mL of YPD at an OD_600_ = 0.2. Cells were grown at 30°C, shaking at 250 rpm, until OD_600_ reached 0.6. Cells were pelleted and resuspended in pre-warmed 37°C YPD. Aliquots of 5mL were pelleted every 30 minutes for 2 hours and flash frozen in liquid nitrogen. Pellets were resuspended in 50 μL of buffer RLT (Qiagen RNeasy kit) + 10μL/mL beta-mercaptoethanol (B-ME), and cells were lysed by mechanical disruption with glass beads using a Bullet Blender. Lysates were resuspended in 450 μL of RLT + B-ME and RNA was extracted using the Qiagen RNeasy kit. For northern blot analysis of *RPL2*, 3 μg of RNA per sample was electrophoretically separated in a 1% agarose + formaldehyde gel. The rRNA was detected and quantified by ImageLab software. RNA was transferred to a nylon membrane, hybridized with a P32-labeled *RPL2* DNA probe, and detected by phosphorimaging using a Typhoon scanner. *RPL2* signal was quantified using ImageLab software. *RPL2* levels were normalized to corresponding rRNA. Statistical analysis was performed using GraphPad Prism (version 8.4.3).

### Western Blot detection of Phospho-Hog1 and Phospho-Mpk1

*C. neoformans* strain *ccr4*Δ and the wild-type parental strain (H99) were used to inoculate 5mL overnight YPD cultures, shaken at 30°C, 250rpm. Overnight cultures were used the following day to inoculate fresh YPD (at OD_600_ = 0.2), followed by 5 hours of shaking in baffled flasks in a 30°C incubator until mid-log phase (OD_600_ = 0.65). Mid-log cells were pelleted and resuspended in pre-warmed YPD and shaken in a 37° incubator at 250rpm, with timepoints collected every half hour. At each time point, cells were pelleted, the supernatant was discarded, and the pellets were rapidly frozen in liquid nitrogen and stored at -80° until lysis. Thawed pellets were rinsed in 1 mL SDW, and then resuspended in whole-cell lysis buffer (15 mM HEPES (pH 7.4), 10 mM KCl, 5 mM MgCl2, 10 μL/mL HALT protease inhibitor, 1mM dithiothreitol (DTT)). Cells were lysed by mechanical disruption using glass beads in a Bullet Blender, and the resulting lysate was clarified by centrifugation. Samples were reduced and denatured by boiling in Laemmli buffer with B-ME and 30 μg of protein per sample was electrophoretically separated through a Bio-Rad Stain-Free Tris-glycine gel. Prior to transferring protein to nitrocellulose, the gel was imaged and total protein was quantified from the gel using ImageLab software. The blot was probed using a rabbit anti-phospho-p38 MAPK (Cell Signaling Technologies), followed by an anti-rabbit HRP secondary antibody and visualized by chemiluminescence using a BioRad ChemiDoc Imager. It was found that the phospho-p38 primary antibody, in addition to detecting phosphorylated Hog1, was cross-reactive with phosphorylated Mpk1. The specificity of this cross-reactive band was verified using a mouse anti-phospho-p42/44 MAPK antibody (Cell Signaling Technologies), which demonstrated loss of the corresponding band in *mpk1*Δ, and reliably showed the same pattern of Mpk1 phosphorylation as the phospho-p38 antibody across all biological replicates. The phospho-p38 signal was selected to show phosphorylation of both Hog1 and Mpk1 due to its higher signal-to-noise ratio.

### RT-qPCR

*C. neoformans* strains *hog1*Δ, *mpk1*Δ, *ccr4*Δ, and the wild-type parental strain (H99) were grown in 5mL overnight cultures in YPD, shaken at 30°, 250rpm. Overnight cμLtures were used the following day to inoculate fresh YPD (at OD_600_ = 0.2), followed by 5 hours of shaking in baffled flasks in a 30° incubator until mid-log phase (OD_600_ = 0.65). Mid-log cells were pelleted and resuspended in pre-warmed YPD and shaken in a 37°C incubator at 250rpm for one hour. Cells were then pelleted, flash frozen in liquid nitrogen, and stored at -80° until further processing. Pellets were thawed on ice and resuspended in buffer RLT (Qiagen RNeasy kit) with 10 μL/mL B-ME, and cells were lysed by mechanical disruption with glass beads using a Bullet Blender. The crude lysate was clarified by centrifugation, and RNA was extracted from the soluble fraction using the Qiagen RNeasy kit. Genomic DNA was digested using the Qiagen RNeasy kit on-column DNase I digestion, and equal quantities of eluted RNA were converted to cDNA using the High-Capacity cDNA Reverse Transcription Kit with RNase Inhibitor (ThermoFisher). qPCR for *CHS4* (CNAG_00546), *CHS5* (CNAG_05818), *CHS6* (CNAG_06487), *EXG1* (CNAG_05803), and mitofusin (CNAG_06688) were performed on cDNA using the qPCR SyGreen Blue Mix LoROX (PCR Biosystems) according to manufacturer instructions. Primers for each gene can be found in Table 1. The data were acquired on a CFX Connect Real-Time PCR Detection System (Bio-Rad). Mitofusin (CNAG_06688) was used as an internal control, and negative control samples without reverse transcriptase were included. Two technical replicates and three biological replicates were performed for all reactions. The ΔΔCt method was used to calcμLate differences in expression. Statistical analysis was performed using GraphPad Prism (version 8.2.1) with significance defined as *\*p<0*.*05, **p<0*.*01, ***p<0*.*001*. One-way ANOVA was performed with Dunnet’s test post hoc comparing the normalized expression of each gene to wild-type H99.

## Results

### A screen of kinase mutants for glucan unmasking implicates core MAP kinases

Reprogramming of the *C. neoformans* translatome through the action of the mRNA deadenylase Ccr4 is required for both temperature adaptation and to maintain masking of b-1,3-glucan at host temperature (Bloom et al., 2019). We reasoned that identification of regulators of glucan masking might also identify factors that control translatome reprogramming in *C. neoformans*. Using b-1,3-glucan unmasking as an output, we screened mutants from the *C. neoformans* kinase deletion collection that were predicted to be sensitive to either host temperature or cell wall stress (Lee et al., 2016), and identified two core mitogen activated protein kinase (MAPK) modules that unmask b-1,3-glucan at 37°C as detected by flow cytometry. Mutants in all three components of the Hog1 MAPK pathway, including the MAP3K *pbs2*D, the MAP2K *ssk2*D and the MAPK *hog1*D exhibit a moderate level of unmasking defined as two-to five-fold over wild type. In addition, all three components of the cell integrity MAPK signaling pathway were found to unmask extensively, defined as unmasking greater than five-fold over wild type. This includes the MAP3K *bck1*D, the MAP2K, *mkk2*D and the MAPK *mpk1*D. We went on to validate the screen in larger-scale cultures, and found that the identified mutants did indeed reproducibly unmask at 37°C (Figure 1). Comparison of glucan exposure by fluorescence microscopy to the WT, which completely masks β-glucan, and the *ccr4*Δ mutant, which exposes β-glucan, confirmed our findings by flow cytometry and revealed extensive unmasking in a speckled pattern around the periphery of a large portion of the cell wall integrity MAPK mutants and moderate unmasking in a crescent-like pattern on the periphery of a subpopulation of the Hog1 MAPK pathway mutants when grown at 37°C (Figure 2).

**Figure 1:**
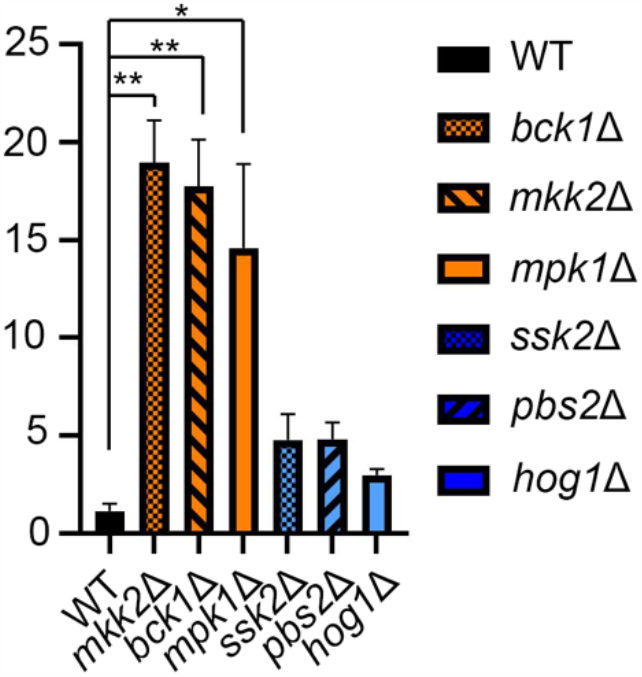
Validation of β-(1,3)-glucan unmasking in kinase mutants of the CWI MAPK and HOG1 MAPK pathways. Kinase mutants grown overnight in YPD at 37°C were stained with anti-β-(1,3)-glucan antibody and assessed for fluorescence by flow cytometry. Fluorescent populations of mutants were compared to WT to calculate fold change in β-(1,3)-glucan exposure. Statistical analysis was performed by Kruskal-Wallis test. Error bars depict SEM, n = 3-4, *\** = *p<0.05*; *\*\** = *p<0.01*.

**Figure 2:**
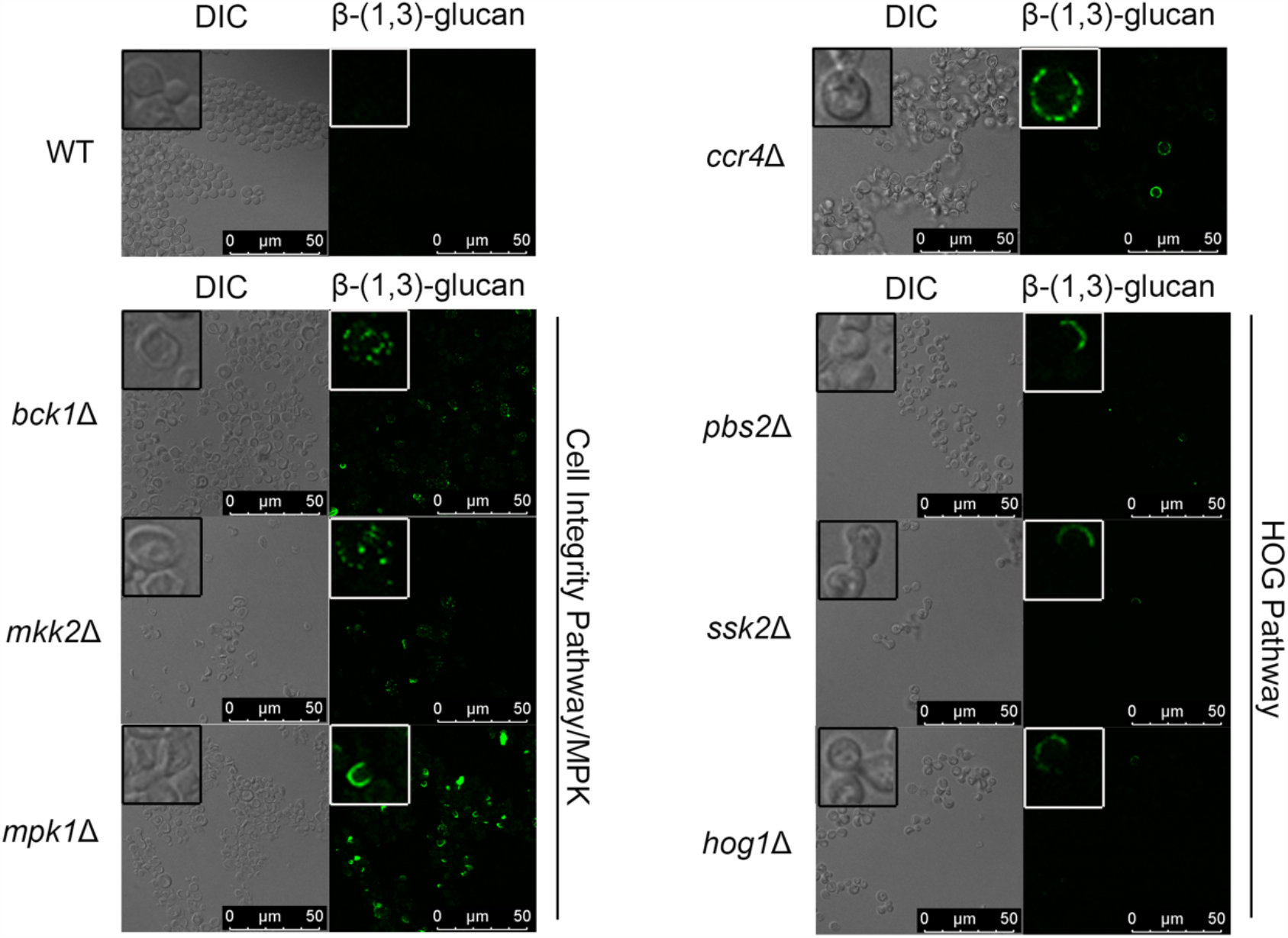
Mutants in the CWI MAPK and HOG MAPK pathways unmask β-(1,3)-glucan at 37C°. Cells grown overnight at 37°C were stained with an anti-β-(1,3)-glucan antibody and imaged by fluorescence microscopy. Insets show a portion of the larger image at 4x magnification.

### Unmasking kinase mutants exhibit differential sensitivity to temperature and cell wall perturbation

Given that the identified kinase mutants exhibit unmasking at host temperature we tested sensitivity to both host temperature stress and cell wall stressors by spot dilution assays (Figure 3A). All of the mutants demonstrated a strong sensitivity to SDS and moderate sensitivity to caffeine. All of the cell integrity MAPK pathway mutants exhibited pronounced sensitivity to congo red, a dye that interacts with β-1,3-glucans, and a modest sensitivity to calcofluor white. None of the mutants exhibited growth defects when spotted onto YPD agar plates and incubated at 37°C, nor were the cell wall stress sensitivities exacerbated at elevated temperature.

**Figure 3:**
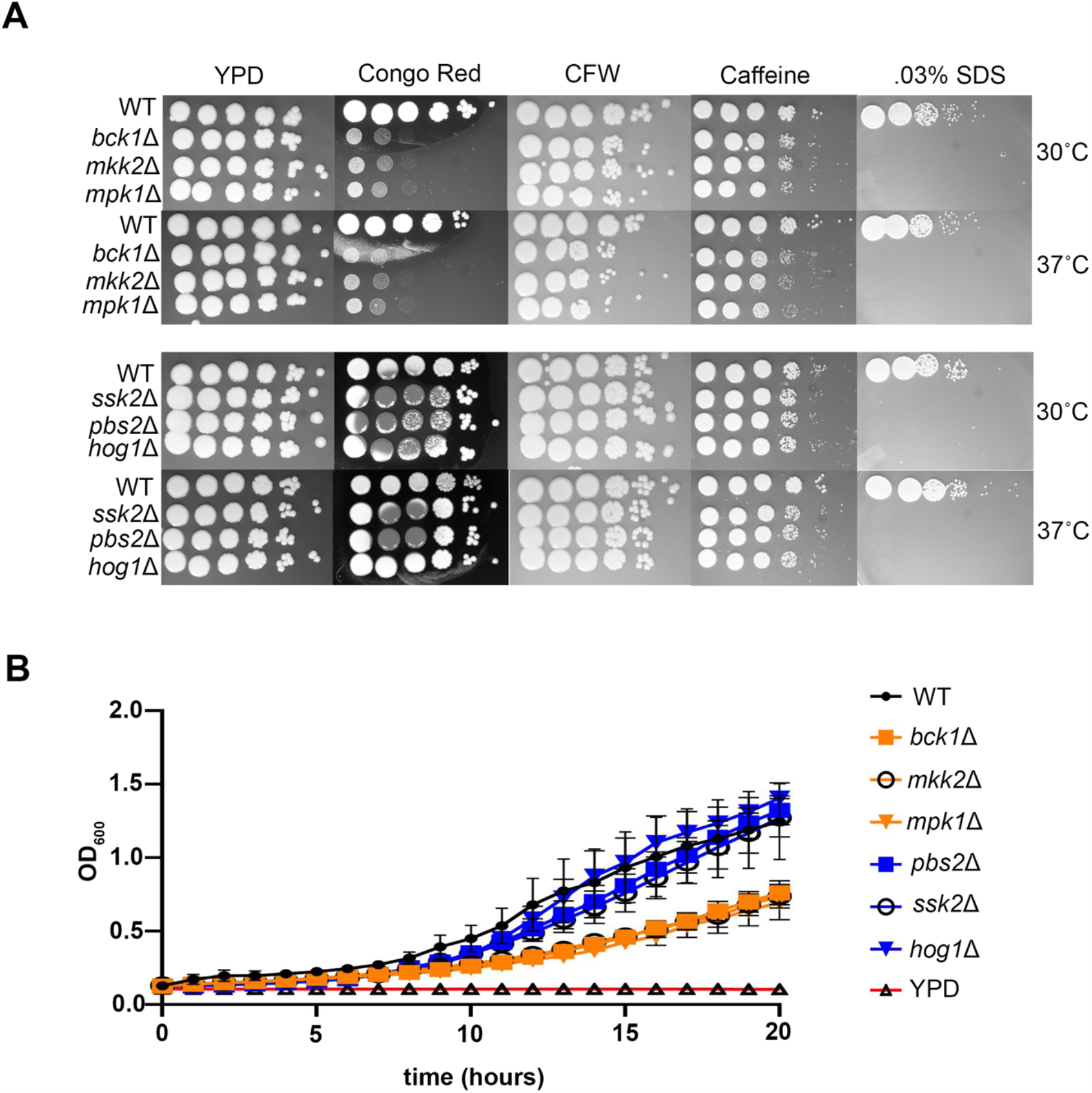
Mutants in the CWI MAPK and HOG MAPK pathways display differential sensitivities to cell wall and host-temperature stress. **(A)** Overnight cultures were diluted to OD_600_ = 1.0, serially diluted 10-fold, spotted on to YPD agar plates containing indicated compounds, and incubated at 30°C or 37°C. **(B)** Mid-logarithmic phase cultures grown at 30°C were diluted to OD_600_ = 0.1 and grown in a 96-well plate at 37°C. OD_600_ was recorded over 20 hours.

To further examine the effect of host temperature we also performed growth curves for the mutants in liquid YPD media at 37°C. We have observed differences in growth between liquid and solid media, and it has recently been shown that humidity plays a role in viability and transmission (Springer et al., 2013). While the Hog1 MAPK mutants demonstrated WT-like growth, all of the CWI MAPK mutants demonstrated slower growth in liquid media despite comparable growth by the spot dilution method (Figure 3B).

### Hog1, but not Mpk1, controls unmasking at the level of translatome reprogramming

Our previous data demonstrate that deadenylation-dependent mRNA decay promotes the reprogramming of the translatome that allows for expression of genes required for temperature adaptation and glucan masking. We next set out to investigate if Hog1 and Mpk1 were acting at the level of translatome reprogramming. The first step in translatome reprogramming is the rapid, but transient, removal of highly abundant and efficiently translated mRNAs from the translating pool (Bloom and Panepinto, 2014). In the wild type, nearly every ribosomal protein mRNA is removed from the translating pool by 1 hour at 37°C (Bloom et al., 2019). Removal of ribosomal protein mRNAs permits the association of transcription factor mRNAs and stress response effector mRNAs with ribosomes. We asked if transient repression of *RPL2*, a representative ribosomal protein mRNA, was intact in the identified unmasking mutants of each pathway. Analysis of *RPL2* repression by northern blot revealed a decreased rate of repression in the *pbs2*D, *ssk2*D, and *hog1*D mutants compared to the wild type (Figure 4). *RPL2* repression in the *bck1*D, *ssk2*D, and *mpk1*D mutants exhibited the same repression kinetics as wild type, with a rapid decline in levels during the first hour followed by a gradual increase toward pre-stressed levels. These data suggest that the Hog1 pathway, but not the Mpk1 cell integrity pathway, regulate the first step in translatome reprogramming.

**Figure 4:**
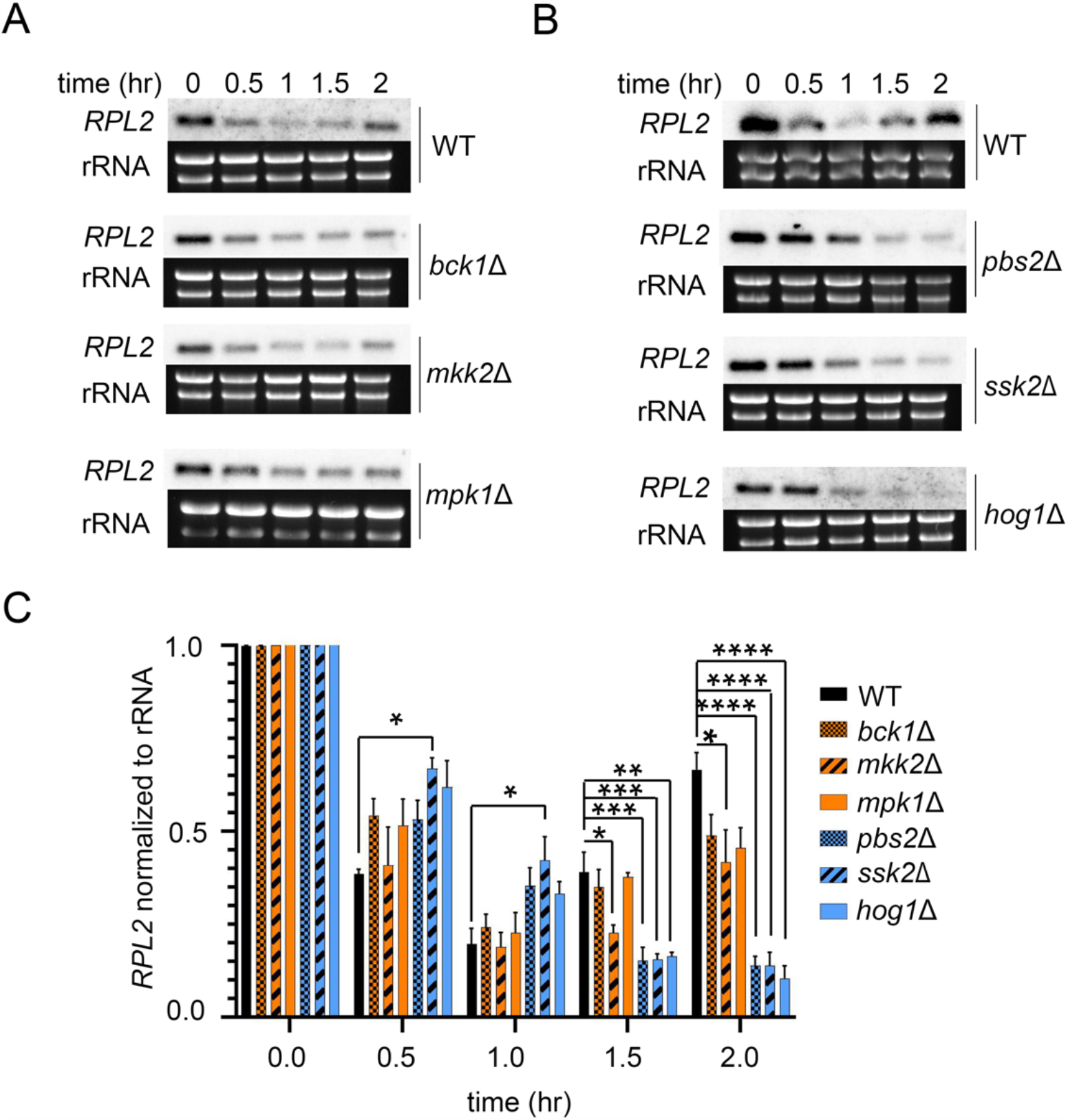
Only Hog1 upstream regulators work at the level of translatome reprogramming. **(A)** Mid-logarithmic cultures grown were shifted to 37°C for 2 hours. *RPL2* levels, normalized to rRNA, were quantified by northern blot analysis. Gels and blots are representative of 3 biological replicates. **(B)** The graph depicts mean *RPL2* levels following a shift to 37°C. Statistical analysis was performed using One-Way ANOVA with Dunnet’s test post hoc, comparing normalized expression of *RPL2* in mutants to WT. Error bars represent the SEM, n=3, *\** = *p<0.05*; *\*\** = *p<0.01, \*\*\** = *p<0.001; \*\*\*\** = *p<0.0001*.

Temperature adaptation in *C. neoformans* is accompanied by a moderate translational repression consistent with the removal of a large group of highly-abundant and highly-translated mRNAs from the translating pool. We reasoned that if RP transcripts were delayed in their removal from the translating pools in the *hog1*D mutant, then we might observe a defect in host-temperature associated changes in the translational landscape. We used polysome profiling to compare the translational state of the wild type, mpk1D and *hog1*D mutant at 30 minutes after a shift to 37°C (Figure 5). Similar to WT, the *mpk1*D mutant exhibited a decrease in low molecular weight polysomes and an increase in high molecular weight polysomes in response to host temperature. The *hog1*D mutant demonstrated a defect in translational regulation with very little change in the polysomes and a large increase the 60s subunit. This is consistent with the Hog1 pathway regulating glucan masking through translatome reprogramming.

**Figure 5:**
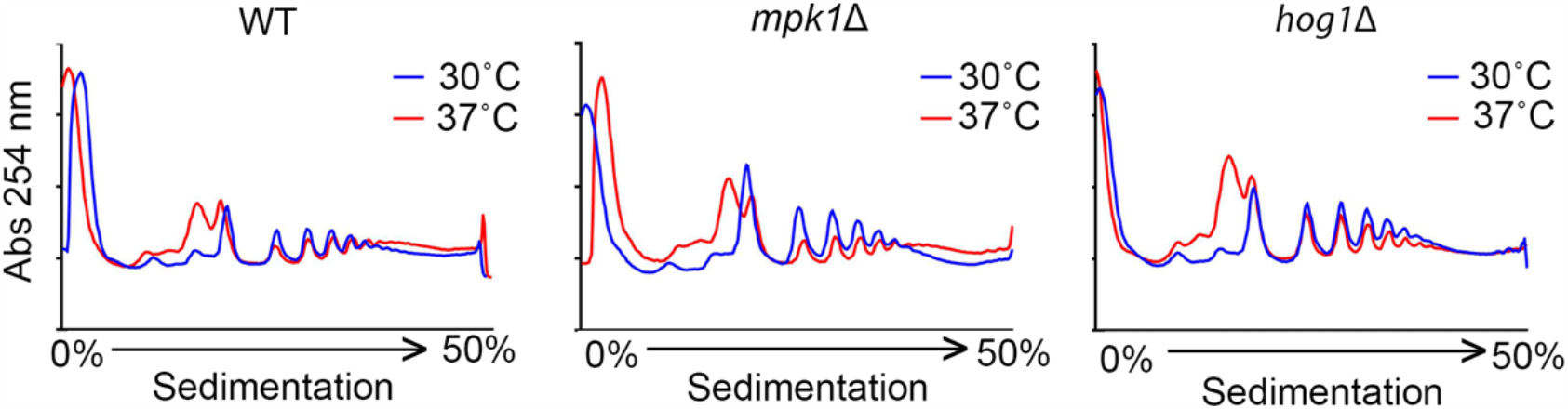
Hog1, but not Mpk1, impacts host temperature induced changes in the translational landscape. Each strain was grown to mid-log phase in YPD at 30°C. Cells were collected from 30°C cultures and cultures that were shifted to 37°C for 30 minutes. Polysome profiles were obtained from reading Absorbance at 254nm of lysates that were separated in sucrose gradients by ultracentrifugation. Profiles are representative of 3 biological replicates.

### Hog1 and Mpk1 are activated during host temperature adaptation

We next sought to establish that these phenotypes in *hog1*Δ and *mpk1*Δ are due to direct participation of these kinase modules during host temperature adaptation. In order to evaluate this, we measured the levels of active phosphorylated Hog1 and Mpk1 by Western blot in WT cells during growth at 37°C (Fig. 6, left). We found that shifting to 37°C activated both kinases in WT, with pronounced activation occurring after 1h.

**Figure 6:**
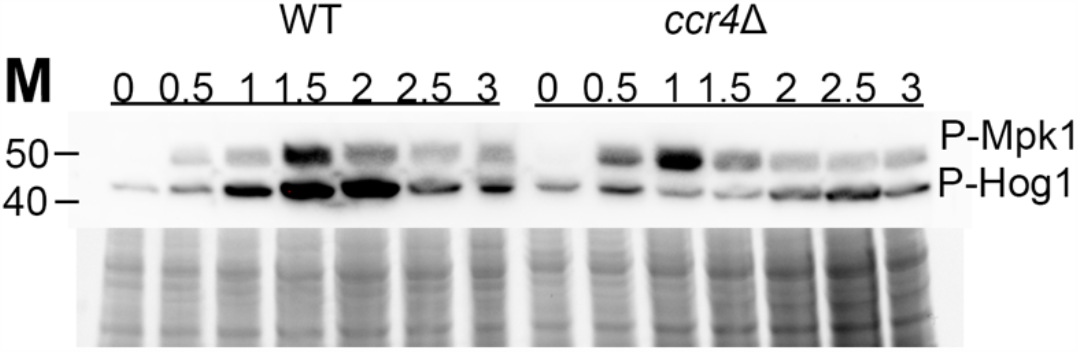
Activation of Hog1 and Mpk1 during growth at mammalian body temperature. WT and *ccr4*Δ cells were grown to midlog phase in YPD at 30°C, then shifted to 37°C for 3 hours, with time points taken every half hour. Lysates were probed by Western blot for phosphorylation of Hog1 and Mpk1 using a cross-reactive antibody. Total protein is shown as a loading control. Data shown are representative of three biological replicates.

The observed delay in RP transcript decay in *hog1*Δ, suggested that Hog1 activity influences removal of RP transcripts from actively translating ribosomes. We wondered then if Hog1 activation would be reduced in the absence of RP transcript decay. To test this, we also examined Hog1 and Mpk1 activation in a *ccr4*Δ mutant, which is severely deficient in RP decay (Bloom et al., 2019) (Fig. 6, right). We found that temperature-induced Hog1 activation was delayed in *ccr4*Δ, beginning after 2 hr, and was not as robust as in wild type. We also observed a small but reproducible acceleration in Mpk1 activation in *ccr4*Δ. Together, these results indicate that the Hog1 and Mpk1 kinase modules directly participate during adaptation to growth at 37°C, and that Ccr4 either directly or indirectly modulates Hog1 activation during host temperature adaptation.

### Ccr4, Hog1 and Mpk1 cooperatively regulate the host-temperature transcriptome

In the absence of deadenylation-dependent mRNA decay, many transcription factors fail to enter the translatome, leading to gross defects in transcriptome reprogramming. In order to evaluate potential genes involved in the glucan masking phenotype, we compared the existing *mpk1*Δ RNA-seq dataset ((Donlin et al., 2014),GSE57217)) to our own dataset from *ccr4*Δ ((Bloom et al., 2019), GSE121183)) (Figure 7A). We filtered the resulting overlap of 232 genes for those that would be cell-surface associated or involved in cell wall remodeling, and among these genes were two chitin synthases, *CHS4* and *CHS5*, as well as an uncharacterized exoglucanase, *EXG1* that were predicted to be down regulated in both sets. We hypothesized that if Hog1 influences removal of RP mRNAs and thus, translatome reprogramming, and if translatome reprogramming precedes transcriptome remodeling as we predict (Figure 8), then these genes would also be affected in the *hog1*Δ mutant. We reported previously that *CHS6* was down regulated in *ccr4*Δ and included that in our analysis. We compared expression of these four genes by RT-qPCR in WT, *mpk1*Δ, *ccr4*Δ, and *hog1*Δ, following a shift to 37°C for one hour (Fig. 7B). We observed a statistically significant downregulation of all four cell wall genes in each of the mutant strains. This finding indicates that Hog1, Mpk1, and Ccr4 all coordinate expression of cell wall genes at 37°C, and suggests that these proteins cooperatively enforce the transcriptional regulon involved in glucan masking.

**Figure 7:**
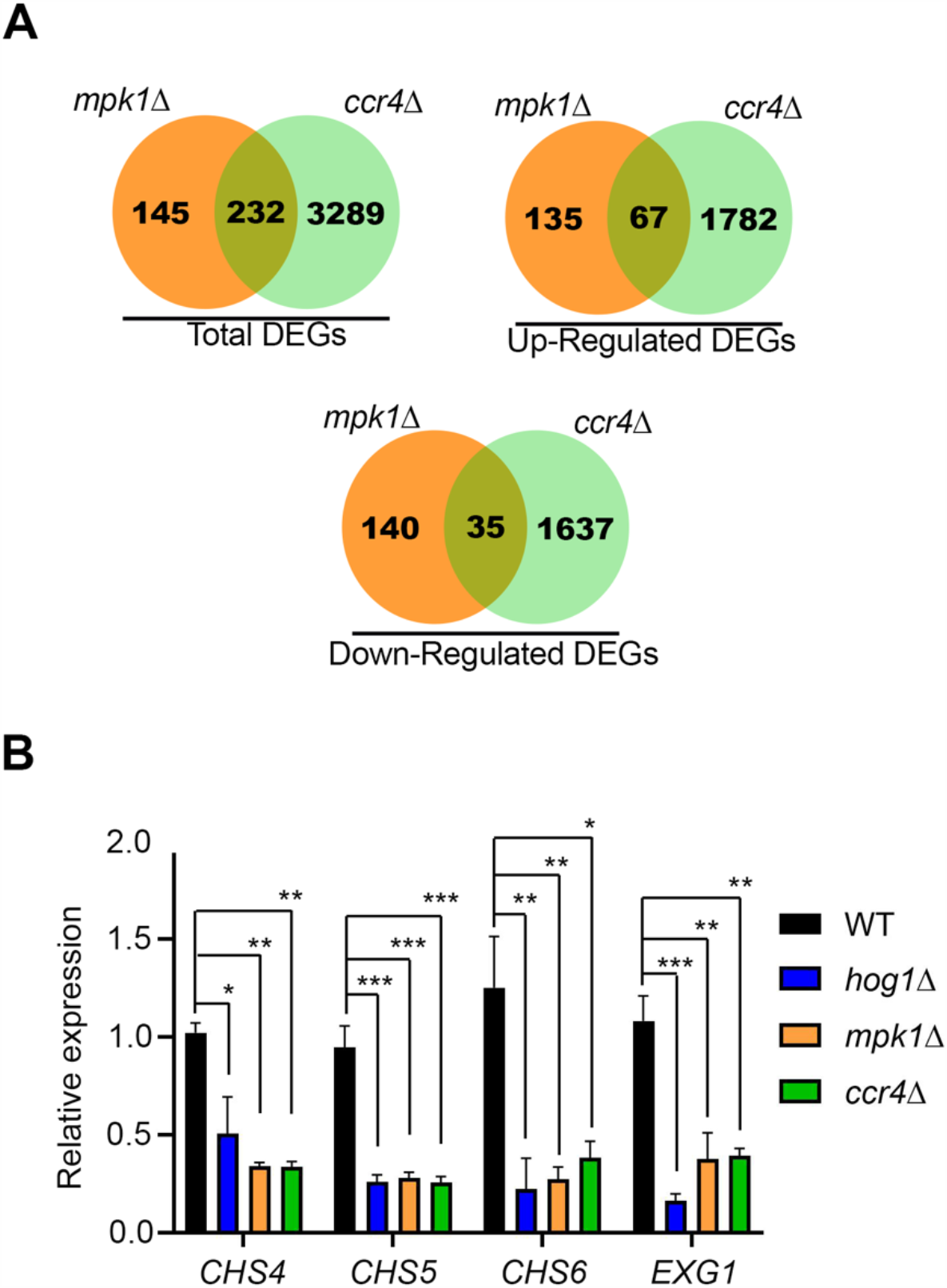
Genes involved in cell wall remodeling are down regulated in unmasking mutants at 37°C. **(A)** Differentially expressed genes in the *mpk1*Δ and *ccr4*Δ strains were compared to identify genes regulated by both the CWI pathway and Ccr4-mediated mRNA decay. **(B)** WT, *hog1*Δ, *mpk1*Δ, and *ccr4*Δ were grown to midlog in YPD at 30°, then shifted to 37° for 1 hr. qPCR was performed on cDNA from these cells to evaluate expression of *CHS4, CHS5, CHS6*, and *EXG1*. The ΔΔCt method was used to calculate expression normalized to mitofusin. Statistical analysis was performed using One-Way ANOVA with Dunnet’s test post hoc, comparing normalized expression of each gene in mutants to WT. Error bars depict SEM, n = 3, *\** = *p<0.05*; *\*\** = *p<0.01*; *\*\*\** = *p<0.001*.

**Figure 8.**
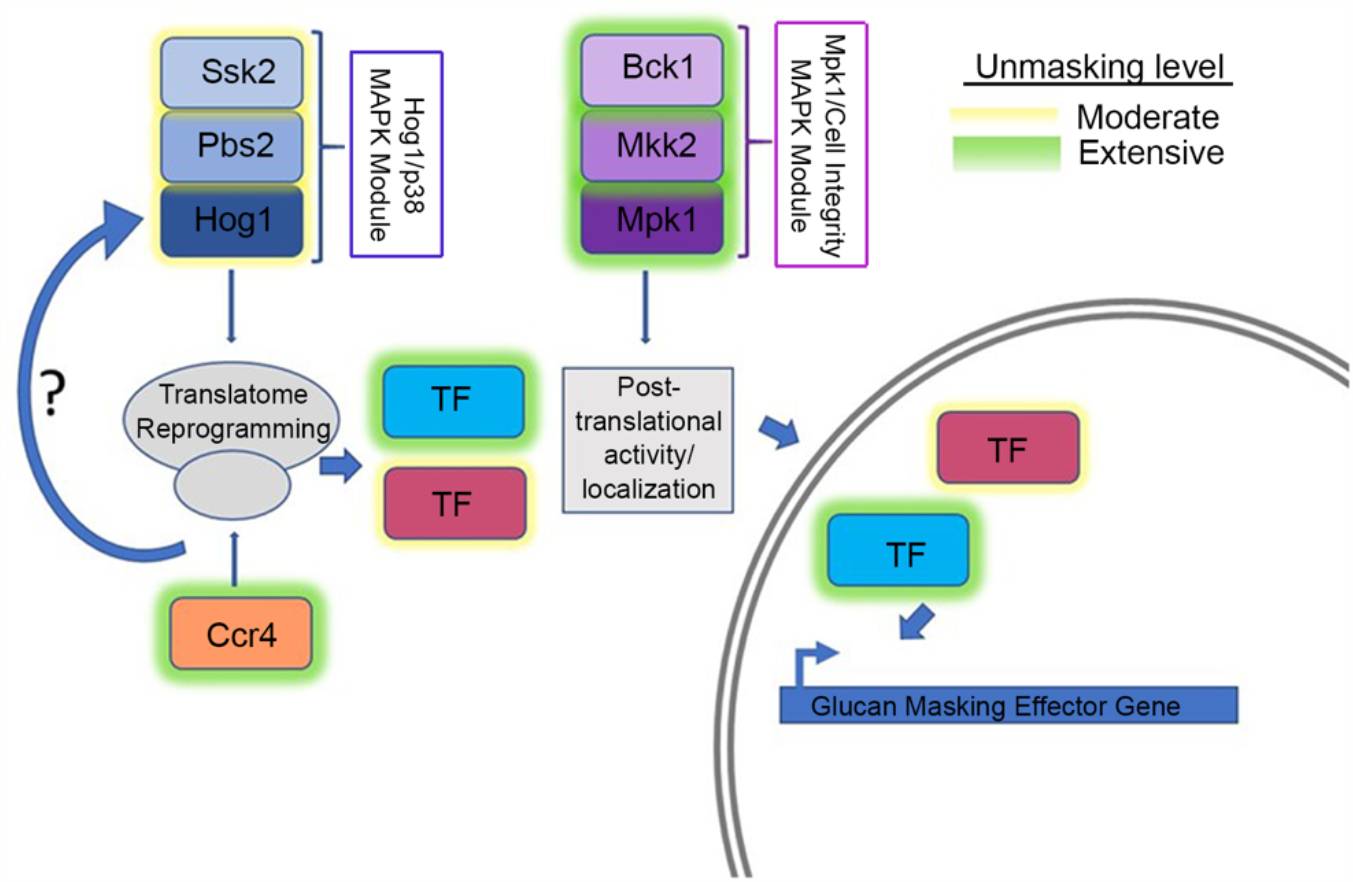
A model for the role of the CWI MPK and HOG MPK pathways in host temperature induced β-(1,3)-glucan unmasking. Our data suggests that the HOG MAPK pathway acts at the level of translatome reprogramming. Decay of RP transcripts may impact the activation of Mpk1, perhaps via a complex feedback loop. The CWI MAPK pathway does not affect translatome reprogramming, but may act downstream on newly synthesized transcription factors that govern glucan masking.

## Discussion

The fungal cell wall is essential and impacts not only the structure and rigidity of the cell, but mediates all interactions the cell encounters with its environment. It is logical then that this structure is dynamic and displays plasticity. β-1,3-glucan is claimed to be the most important component of fungal cell walls, serving to anchor other components by covalent attachment (Garcia-Rubio et al., 2019). It is also highly antigenic in the context of a host, and exposure of this glucan results in rapid detection by Dectin-1 resulting in protective immune responses (Taylor et al., 2007). Pathogenic fungi utilize mechanisms of glucan masking to help evade the host immune system. While *C. albicans* hides its glucan layer under a blanket of mannoproteins (Gantner et al., 2005), *Asperigullus* species mask β-1,3-glucan with the hydrophobic surface protein, RodA on conidia (Carrion Sde et al., 2013) and a polysaccharide cloak of galactosaminogalactan on hyphae (Gravelat et al., 2013), and *C. neoformans* is protected by a polysaccharide capsule (O’Meara and Alspaugh, 2012). The continual changes that a pathogen encounters in a host must elicit responses that include cell wall remodeling to keep it cloaked from attack.

Unlike other pathogenic fungi, β-1,3-glucan is not the most abundant cell wall carbohydrate in *C. neoformans*, however, it is essential as deletion of the sole β-1,3-glucan synthase, *FKS1*, is lethal (Thompson et al., 1999; Gilbert et al., 2010). Studies investigating β-1,3-glucan masking have been lacking in this organism. We previously reported that Ccr4-mediated mRNA decay and translatome reprogramming in response to host temperature was required for keeping β-1,3-glucan masked. Here, we identified additional signaling modules that mediate this outcome.

The Pkc1 CWI MAPK pathway is the major pathway that regulates cell wall integrity in *C. neoformans* (Gerik et al., 2005), thus, we were not surprised to find that mutants in this pathway unmasked β-1,3-glucan. Like the Cek1 MAPK pathway in *C. albicans* (Galan-Diez et al., 2010), this pathway may control masking in a general manner by regulating genes that are involved in cell wall structure even under no stress as these mutants are inherently sensitive to cell wall stressors. In fact, mutation of the most upstream kinase of this pathway, Pck1, is only viable in the presence of an osmostabilant (Gerik et al., 2008). Host temperature may increase reliance on this pathway to help in combating stress-induced insult to the cell wall. The terminal kinase, Mpk1 was previously reported to be activated in response to oxidative and nitrosative stresses (Gerik et al., 2008), and we show here that it is activated in response to 37°C and may explain both the growth defect in mutants in this pathway when grown in liquid culture at 37°C and the extensive β-(1,3)-glucan unmasking (Figures 1, 3B). We showed that Mpk1 activation in response to temperature stress was not hindered when translatome reprogramming is inhibited by defective clearance of ribosomes in the *ccr4*Δ mutant (Figure 6). Further, RP transcript repression (Figure 4) and polysome profiles (Figure 5) were unaffected in the *mpk1*Δ mutant suggesting that the CWI MAPK pathway does not impact unmasking at the level of translatome reprogramming. Given that our previous data suggests that translatome reprogramming impacts transcription factor production and precedes transcriptional remodeling, we speculate that CWI MPK regulation of masking in response to host temperature acts in parallel with or after translatome reprogramming, perhaps through post-translational modification of newly translated transcription factors and thereby affecting their activity or localization (Figure 8). Indeed, our previous work identified several genes that encode transcription factors that are down regulated in the *ccr4*Δ mutant that may be influenced by Mpk1, which was supported by the overlapping differentially expressed genes in these two strains (Figure 6A). Importantly, we note that the RNAseq data for the *mpk1*Δ strain was acquired from cells grown in YPD supplemented with the osmostabilant sorbitol due to the original inclusion of the *pck1*Δ mutant in those studies (Donlin et al., 2014).

The *C. neoformans* HOG MPK pathway has been studied largely in regard to osmoregulation and drug resistance (Bahn et al., 2005; Ko et al., 2009; Kim et al., 2011). Interestingly, Hog1 negatively regulates capsule production, and in response to osmotic stress the constitutive activation of Hog1 in *C. neoformans* is down regulated (Bahn et al., 2005; Ko et al., 2009). We show here that Hog1 activation is heightened in response to host temperature, and this activation is dependent, at least in part, on Ccr4-mediated clearance of actively translating ribosomes (Figure 6). Intriguingly, the absence of *HOG1* resulted in significantly delayed repression of the abundant RP transcripts following a shift to 37°C. These mRNAs need to be cleared from ribosomes for translatome reprogramming to ensue. Delayed RP repression was supported by the lack of change in the translational landscape in response to host temperature in the *hog1*Δ mutant. If Hog1 activation was a prerequisite for Ccr4-mediated decay, then we would have expected that Hog1 phosphorylation would occur in the *ccr4*Δ mutant similarly to WT, but this was not the case. The relationship between Hog1 activation and translatome reprogramming is therefore complex, and it begs to be answered at which time point in temperature response Hog1 is acting. In *Saccharomyces cerevisiae*, Hog1 has been indirectly linked to translational regulation via activation of its downstream target Rck2, which targets the translation elongation factor eEF2 (Teige et al., 2001). We are currently investigating the events that occur at the ribosome in response to stress to identify the order of operations that result in translatome reprogramming. The data revealed here suggest that Hog1 is required for the immediate repression of RP transcripts and their exit from the translating pool to allow for translation of newly synthesized genes such as those involved in masking (Figure 8). Given that the *hog1*Δ mutant grows as well as WT at 37°C suggests that translatome reprogramming eventually occurs, however glucan masking is not completely achieved. Future assessment of phagocytosis will determine if Hog1-mediated β-(1, 3)-glucan masking contributes to host evasion and if loss of Hog1-mediated β-(1, 3)-glucan masking alters the immune landscape in the lung.

Our work here is the first to examine signaling modules that specifically play a role in masking of the antigenic cell wall component, β-(1,3)-glucan in *C. neoformans*. Given that innate immunity is equipped to specifically recognize this pattern associated molecular pattern and elicit protective immune responses, the understanding of the pathogen’s ability to cloak itself may be essential in future therapeutic endeavors. That protein translation has proven to be targetable, the role of factors that specifically manipulate the translational machinery to promote pathogen adaptation could be instrumental.

## Author Contributions

AB, DG, and JP conceived and designed the experiments. AB, DG, and CK performed the experiments. AB, DG, and JP wrote the manuscript.

## Acknowledgements

The kinase deletion collection was obtained from the Fungal Genetics Stock Center (Manhattan, Kansas, USA).

